# Multiscale modeling of HBV infection integrating intra- and intercellular viral propagation for analyzing extracellular viral markers

**DOI:** 10.1101/2023.06.06.543822

**Authors:** Kosaku Kitagawa, Kwang Su Kim, Masashi Iwamoto, Sanae Hayashi, Hyeongki Park, Takara Nishiyama, Naotoshi Nakamura, Yasuhisa Fujita, Shinji Nakaoka, Kazuyuki Aihara, Alan S. Perelson, Lena Allweiss, Maura Dandri, Koichi Watashi, Yasuhito Tanaka, Shingo Iwami

## Abstract

Chronic infection of hepatitis B virus (HBV) is caused by the persistence of closed circular DNA (cccDNA) in the nucleus of infected hepatocytes. Despite available therapeutic anti-HBV agents, eliminating the cccDNA remains challenging. The quantifying and understanding dynamics of cccDNA are essential for developing effective treatment strategies and new drugs. However, it requires a liver biopsy to measure the intrahepatic cccDNA, which is basically not accepted because of the ethical aspect. We here aimed to develop a non-invasive method for quantifying cccDNA in the liver using surrogate markers present in peripheral blood. We constructed a multiscale mathematical model that explicitly incorporates both intracellular and intercellular HBV infection processes. The model, based on age-structured partial differential equations (PDEs), integrates experimental data from in vitro and in vivo investigations. By applying this model, we successfully predicted the amount and dynamics of intrahepatic cccDNA using specific viral markers in serum samples, including HBV DNA, HBsAg, HBeAg, and HBcrAg. Our study represents a significant step towards advancing the understanding of chronic HBV infection. The non-invasive quantification of cccDNA using our proposed methodology holds promise for improving clinical analyses and treatment strategies. By comprehensively describing the interactions of all components involved in HBV infection, our multiscale mathematical model provides a valuable framework for further research and the development of targeted interventions.

## Introduction

Chronic hepatitis B virus (HBV) infection is a major public health problem, affecting approximately 297 million people worldwide (https://www.who.int/en/news-room/fact-sheets/detail/hepatitis-b), and increasing the likelihood of developing hepatocellular carcinoma. According to the World Health Organization (WHO) report, HBV infection caused the death of 820,000 people in 2019. Currently, Pegylated interferon alpha (PEG IFN-α) and nucleos(t)ide analogues (NAs) are used as therapeutic agents for chronic hepatitis B (CHB) [1]. PEG IFN-α suppresses viral replication by activating the host’s immune response, while nucleoside analogs strongly reduce the amount of HBV DNA by inhibiting reverse transcription [2]. These treatments are effective in reducing viral load and thereby improving hepatitis, but they are not curative, largely due to the persistence of closed circular DNA (cccDNA), which is responsible for CHB, in the patients in many cases [3]. Therefore, understanding the amount and dynamics of cccDNA is crucial for the development of effective therapeutic strategies. To assess the eradication of cccDNA, liver biopsy is typically required to quantify its amount, although this procedure is not commonly performed in clinical practice.

HBV infects host hepatocytes via binding to the viral receptor, sodium-taurocholate co-transporting polypeptide (NTCP), and is then transported to the nucleus to form cccDNA [4]. The cccDNA is a template for viral replication, which produces viral mRNAs with different lengths. One of the transcripts with approximately 3.5 kb in length, called pre-genomic (pg) RNA is reverse transcribed into HBV DNA. Additionally, the cccDNA stimulates the production of viral proteins such as HBV surface antigen (HBsAg) and HBV core-related antigen (HBcrAg). Integrated HBV DNA (iDNA) of host chromosomes also contributes to the presence of HBV antigens, particularly HBsAg [5, 6]. HBsAg forms the viral envelope and is released to the serum as either an infectious particle with HBV DNA or a subviral particle. HBcrAg includes HBV core antigen (HBcAg) which forms a viral capsid, in addition to HBV e antigen (HBeAg) and a truncated core-related protein called p22cr as nonstructural viral proteins [7, 8]. Thus, cccDNA in hepatocytes plays a crucial role in the persistence of HBV infection.

Mathematical modeling plays a crucial role in understanding the complex dynamics of viral infections [9-11]. In the case of hepatitis B virus (HBV) infection, mathematical models described by ordinary differential equations (ODEs) have been extensively used to investigate various aspects of the infection process [12, 13]. By capturing the dynamics of intercellular HBV infection, these mathematical models have enabled us to quantify the reduction of HBV DNA during therapy [14-16]. The decline of HBV DNA can be accurately estimated by mimicking the dynamics of intracellular HBV replication as ODEs [12, 13, 17]. Moreover, a mathematical model has been employed to explore the role of HBsAg production from integrated DNA in the process of HBV infection [13]. Utilizing clinical data, intercellular infection models have been able to distinguish between the kinetics of the noncytolytic and cytolytic immune responses during acute HBV infection [18]. While mathematical models have been proposed to investigate the impact of HBeAg on inducing immunological tolerance during HBV infection [19] and have suggested the inclusion of antibody response [20, 21], further mathematical models are needed to elucidate the immune response [22]. In contrast to the multiscale models described by partial differential equations (PDEs) developed to study the effects of drugs on HCV infection, there is a notable absence of multiscale models for HBV infection that integrate intracellular and intercellular dynamics [23]. Furthermore, since the primary goal of therapy currently is to achieve functional cure with cccDNA inactivation, very few studies have incorporated cccDNA, the molecular reservoir of HBV, into mathematical models and linked it to experimental data [17, 22].

In this study, our aim was to devise a non-invasive method for quantifying intrahepatic cccDNA in vivo, employing surrogate markers present in peripheral blood. To achieve this goal, by taking an advantage of describing the interactions of all components, we developed a multiscale mathematical model that explicitly incorporates both intracellular and intercellular HBV infection processes (e.g., [24]). Based on age-structured PDEs, our model enables the quantification of HBV viral dynamics, utilizing experimental data gathered from in vitro and in vivo investigations. By applying our multiscale mathematical model, we successfully predicted the amount and dynamics of intrahepatic cccDNA. This was accomplished by quantifying specific viral markers, namely HBV DNA, HBsAg, HBeAg, and HBcrAg, in serum samples. This methodology holds promise for advancing our understanding of chronic HBV infection and may pave the way for improved clinical analyses and treatment strategies.

## Results

To predict the amount of intrahepatic cccDNA under antiviral treatment, we developed a multiscale mathematical model that explains the process of intracellular and intercellular HBV propagation using data from cell culture experiments and humanized mice models in a multiscale iterative model construction manner. First, we used cell culture experiments to create a minimum mathematical model of intracellular HBV replication with or without antivirals, which allowed us to measure cccDNA in hepatocytes over time (**Fig. 1**). Next, we developed a multiscale model of intracellular and intercellular HBV infection *in vivo*, integrating the minimum model into a standard intercellular virus infection dynamics model. We used the humanized mice model to evaluate the performance of this model by measuring longitudinal extracellular viral markers (e.g., HBV DNA, HBsAg, HBcrAg, and HBeAg) in peripheral blood and cccDNA levels in hepatocytes from sacrificed mice before and after treatment (**Fig. 2**). We explain our multi-experimental integrated modeling approach in more detail below, highlighting the link between these two models.

**Figure 1.**
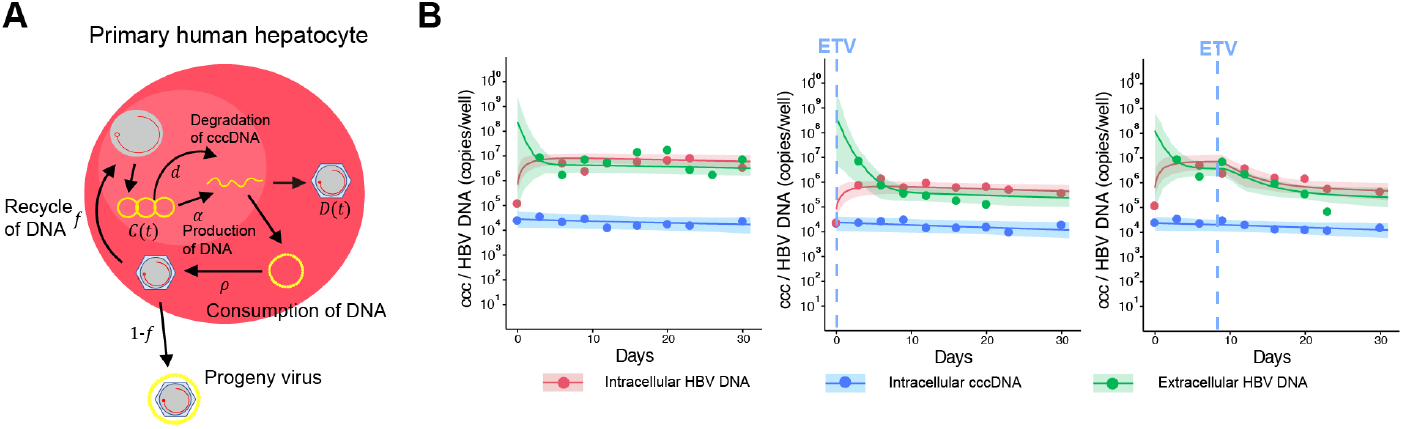
Dynamics of HBV infection in PHH cells: **(A)** Modeling of the intracellular viral life cycle in HBV-infected primary human hepatocytes is shown. Intracellular HBV DNA is produced from cccDNA at rate *α* and is consumed at rate ρ ρ. That is, a fraction 1 − *f* of HBV DNA assembled with viral proteins as virus particles is exported from infected cells, and the other fraction *f* is reused for further cccDNA formation having a degradation rate of *d*. **(B)** Fits of the mathematical model (solid lines) to the experimental data (filled circles) on intracellular HBV DNA and cccDNA, and extracellular HBV DNA in PHH without treatment, or treated with ETV at different times post-infection (red: intracellular HBV DNA, blue: intracellular cccDNA, green: extracellular HBV DNA). The shaded regions correspond to 95% posterior intervals and the solid curves give the best-fit solution (mean) for Eqs. (1-4) to the time-course dataset. All data were fitted simultaneously.

**Figure 2.**
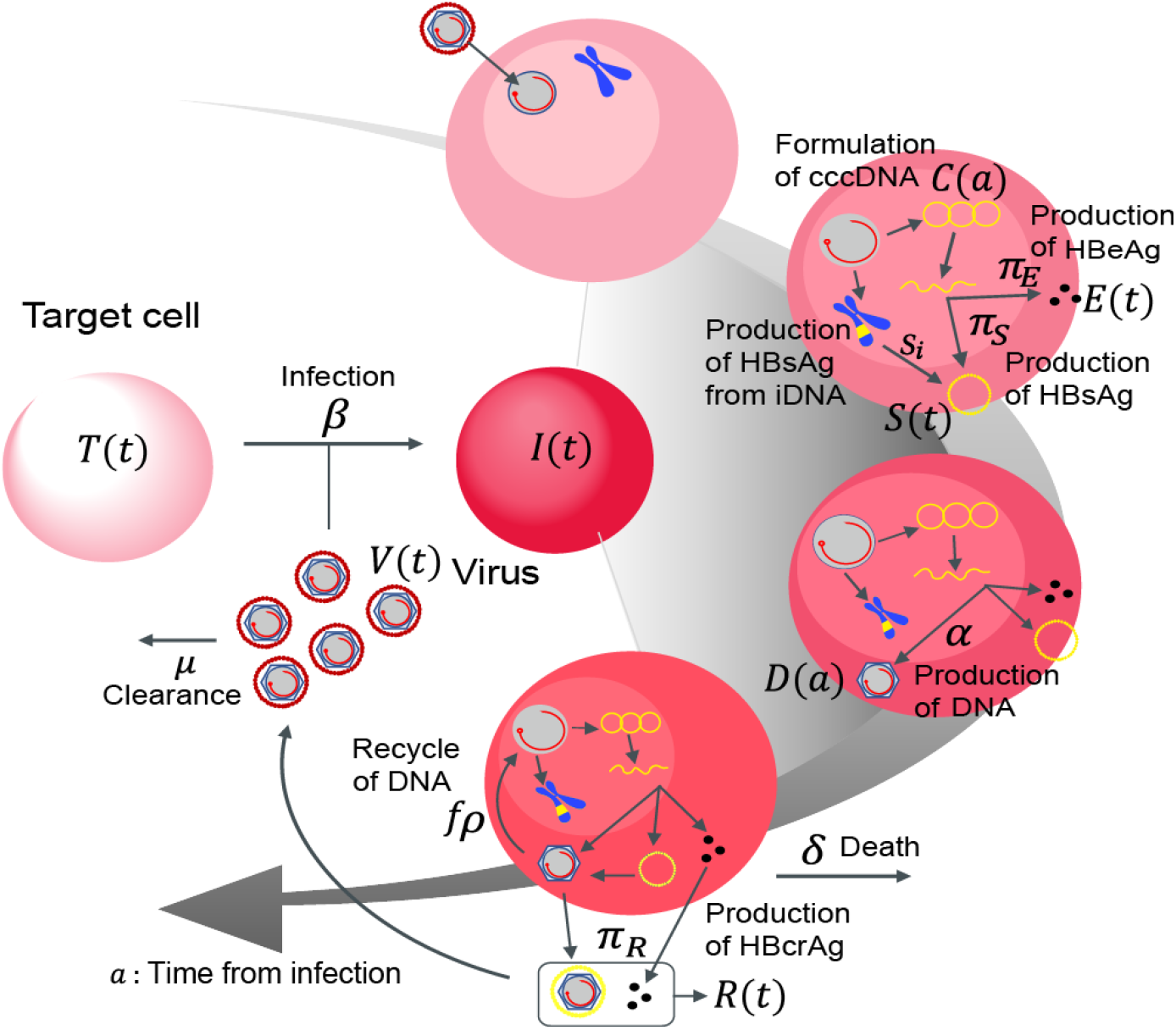
Schematic diagram of multiscale modeling of intracellular and intercellular infection: The entry virion forms cccDNA in the nucleus and produces intracellular HBV DNA at rate *α*. HBsAg, HBeAg, and HBcrAg antigens are also produced from cccDNA at rates *π*_*S*_, *π*_*E*_, and *π*_*R*_ and cleared at *σ* in peripheral blood, respectively. Additionally, HBsAg may also be produced by integrated HBV DNA (iDNA) in the infected cells at a rate *s*_*i*_. The intracellular HBV DNA is consumed at rate ρ ρ, of which a fraction 1 − *f* of HBV DNA assembled with viral proteins as virus particles is exported from infected cells and the other fraction *f* is reused for further cccDNA formation having a degradation rate of *d*. The infected cells are dead at rate *δ* and the exported viral particles, which are cleared at rate *μ*, infect their target cells at rate *β*.

### Mathematical model of intracellular HBV replication dynamics: Intracellular data by a cell culture model

To develop a minimum mathematical model reflecting the dynamics of HBV propagation including cccDNA, we performed cell culture experiments using primary human hepatocytes (PHH) because cccDNA can be “directly” quantified in this system (**Fig. 1A, Fig. S1A** and **Methods**). PHH were infected with HBV and the amount of extracellular and intracellular HBV DNA and intracellular cccDNA were monitored longitudinally (every three to four days up to 24-31 days post-inoculation) either with or without drug treatment (**Fig. 1B, Fig. S1A** and **Methods**). Note that PHH were maintained at 100% confluent conditions with a 2% concentration of dimethyl sulfoxide (DMSO) in the medium during the entire infection assay to support low cell growth and prevent cell division [25-27].

To describe the intracellular virus life cycle in HBV-infected PHH, we developed the following mathematical model (**Fig. 1A**):

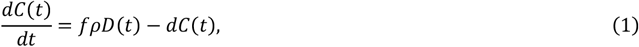

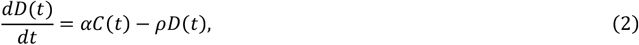

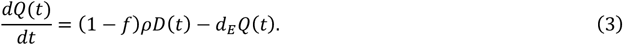

The variables *C*(*t*), *f*(*t*) and *Q*(*t*) represent the amount of intracellular cccDNA and intracellular and extracellular HBV DNA in cultures that have been infected for time *t* (later, we consider infection age as *a* instead of *t*), respectively. The intracellular HBV DNA is produced from cccDNA at rate *α* and is lost at rate ρ ρ of which a fraction 1 − *f* of HBV DNA is assembled with viral proteins as virus particles that are exported out of infected cells, and the other fraction *f* is reused for further cccDNA formation. The viral particles have a clearance rate *d*_*E*_ and cccDNA has a degradation rate of *d*. We have ignored the degradation of intracellular DNA since it is small compared with the consumption rate of HBV DNA due to virion production [28, 29] (see **Table 1**). This intracellular HBV replication model can be modified to include the antiviral effects of different classes of drugs. For example, under treatment with entecavir (ETV), which is a reverse transcriptase inhibitor, the antiviral effect of ETV is assumed to be in blocking HBV DNA production with an effectiveness, *ε*, 0 < *ε* ≤ 1, and is modelled by assuming

**Table 1.** Estimated parameters and initial values for HBV infection in PHH.

| Parameter or variable | Symbol | Unit | Mean | 95% CI |
| --- | --- | --- | --- | --- |
| Production rate of HBV DNA from cccDNA | $\alpha$ | day<sup>-1</sup> | $2.14 \times 10^2$ | $(0.62 - 6.32) \times 10^2$ |
| Fraction of HBV DNA recycling for cccDNA | $f$ | --- | $1.26 \times 10^{-5}$ | $2.71 \times 10^{-10} - 1.38 \times 10^{-4}$ |
| Degradation rate of cccDNA | $d$ | day<sup>-1</sup> | $1.90 \times 10^{-2}$ | $(0.34 - 4.58) \times 10^{-2}$ |
| Consumption rate of HBV DNA for virion | $\rho$ | day<sup>-1</sup> | $6.49 \times 10^{-1}$ | 0.21 – 1.77 |
| Clearance rate of extracellular HBV DNA | $d_E$ | day<sup>-1</sup> | 1.10 | 0.45 – 2.47 |
| Inhibition rate of ETV | $\varepsilon$ | --- | 0.89 | 0.75 – 0.97 |
| Initial value for cccDNA * | $C(0)$ | copies/well | $2.87 \times 10^4$ | $(1.38 - 5.23) \times 10^4$ |
| Initial value for cccDNA ** | $C(0)$ | copies/well | $2.31 \times 10^4$ | $(1.21 - 4.03) \times 10^4$ |
| Initial value for cccDNA *** | $C(0)$ | copies/well | $2.36 \times 10^4$ | $(1.25 - 403) \times 10^4$ |
| Initial value for intracellular HBV DNA* | $D(0)$ | copies/well | $1.89 \times 10^5$ | $(1.75 - 7.90) \times 10^5$ |
| Initial value for intracellular HBV DNA ** | $D(0)$ | copies/well | $3.52 \times 10^5$ | $(0.03 - 1.46) \times 10^5$ |
| Initial value for intracellular HBV DNA *** | $D(0)$ | copies/well | $1.76 \times 10^5$ | $(1.80 - 7.58) \times 10^5$ |
| Initial value for extracellular HBV DNA * | $Q(0)$ | copies/well | $1.53 \times 10^8$ | $1.61 \times 10^4 - 1.35 \times 10^9$ |
| Initial value for extracellular HBV DNA ** | $Q(0)$ | copies/well | $3.63 \times 10^8$ | $2.88 \times 10^4 - 3.82 \times 10^9$ |
| Initial value for extracellular HBV DNA *** | $Q(0)$ | copies/well | $1.80 \times 10^8$ | $1.66 \times 10^4 - 1.62 \times 10^9$ |

\* These values are estimated for condition 1. \*\* These values are estimated for condition 2. \*\*\* These values are estimated for condition 3.

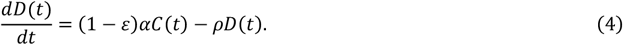

Then we fitted the model to the time-course quantification datasets obtained with and without treatment with ETV (**Methods**). The inhibition of HBV DNA production by ETV perturbates intracellular HBV replication, which enabled us to estimate parameters in the mathematical model [9]. In fact, the amounts of intracellular and extracellular HBV DNA are decreased after treatment with ETV (the middle and right panels in **Fig. 1B**) compared with the control experiment (the left panel in **Fig. 1B**), whereas the amount of intracellular cccDNA is not changed because of the different time scales of decay for HBV DNA and cccDNA (see **Table 1**). The typical behaviour of the model using these best-fit parameter estimates is shown together with the data in **Fig. 1B**, and the estimated parameters and initial values are listed in **Table 1**. It was estimated that 214 copies of HBV DNA are produced from cccDNA in a cell per day on average; only 0.00126% of the produced HBV DNA is recycled to cccDNA. The mean half-life of cccDNA is 51 days in PHH, which is consistent with previous results showing the cccDNA half-life and the limited recycling activity in PHH [26, 27, 30].

### Multiscale mathematical model of intracellular and intercellular HBV infection dynamics

While we can “directly” monitor cccDNA dynamics in hepatocyte cell culture experiments (**Fig. 1B**), it is difficult to obtain time-course measurements of cccDNA *in vivo*. Next, we thus extended the above combined experimental-theoretical approach to describe HBV dynamics *in vivo* and to estimate the cccDNA half-life using extracellular viral markers present in peripheral blood. Here, to precisely quantify both intracellular and intercellular virus dynamics from these viral markers, we introduce a multiscale model using partial differential equations (PDEs) that couple intra-, inter- and extra-cellular virus dynamics for analyzing multiscale experimental data of HBV infection (c.f. [10]) (**Fig. 2**):

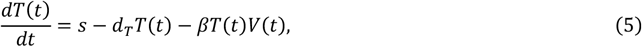

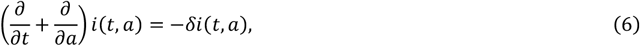

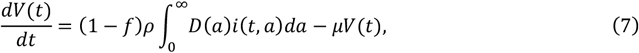

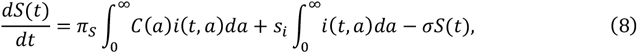

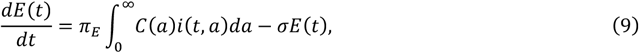

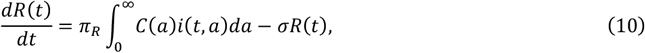

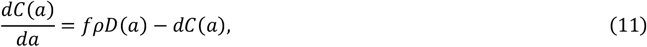

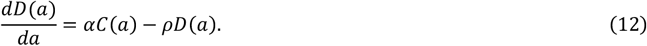

with the boundary condition *i*(*t*, 0) = *βT*(*t*)*V*(*t*) and initial condition *i*(0, *a*) = *i*_0_(*a*). The intercellular variables *T*(*t*) and *V*(*t*) are the number of uninfected cells and the (extracellular) HBV DNA load, respectively. We defined the density of infected cells with infection age *a* as *i*(*t, a*), and therefore the total number of infected cells is 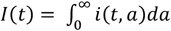. The intracellular variables *C*(*a*) and *D*(*a*), which evolve depending on the age *a*, represent the amount of intracellular cccDNA and HBV DNA, respectively. We also defined extracellular variables used as “extracellular viral markers” to predict the dynamics of cccDNA in hepatocytes, that is, the amount of HBsAg, HBeAg and HBcrAg antigens as *S*(*t*), *E*(*t*) and *R*(*t*), respectively. The definition of an age-structured population model is found in [31]. In addition to the intracellular HBV replication dynamics (see **Fig. 1A**), we assumed target cells, *d*, are supplied at rate *s*, die at per capita rate *d*_*T*_, are infected by viruses at rate *β*, and the infected cells die at per capita rate *δ*. We also considered that HBsAg, HBeAg and HBcrAg antigens are produced from cccDNA in infected cells at rates *π*_*S*_, *π*_*E*_ and *π*_*R*_, and are cleared at rate *σ*, respectively. Additionally, HBsAg may also be produced by integrated HBV DNA (iDNA) in the infected cells at a rate *s*_*i*_. The exported viral particles, i.e., extracellular HBV DNA load, is assumed to be cleared at rate *μ* per virion.

### Analyzing extracellular viral markers by a humanized mice model

To check the performance of our multiscale model, we conducted an HBV infection experiment with humanized liver uPA/SCID mice: after mice were inoculated with HBV and a sustained HBV DNA load was reached (approximately 5.6 × 10^8^ copies/ml) at 53 days post-inoculation, mice were treated with or without ETV or PEG IFN-α continuously. We then longitudinally monitored four different viral markers in the peripheral blood every three to seven days up to 70 days post-treatment: extracellular HBV DNA, HBcrAg, HBeAg, and HBsAg (**Fig. S1B** and **Methods**).

We used the multiscale mathematical model of HBV infection (Eqs.(5-12)), in which an infected cell produces progeny HBVs extracellularly that are then degraded or infect other cells. We derived simple linearized equations (Eqs.(S17-S20) and Eqs.(S30-S33)) for fitting to the time-course datasets quantified with mice with or without ETV or PEG IFN-α treatment (**Supplementary Note 1**,**2**,**3**). Here we assumed the proportion of HBsAg produced from iDNA (i.e., *x* in **Supplementary Note 2**,**3**) is fixed to be 0 (all HBsAg is from cccDNA), 0.5 (HBsAg is equally from cccDNA and iDNA) or 0.8 (HBsAg is dominantly from iDNA) as our sensitivity analysis. Note that the decay rates of infected cells were estimated separately from human albumin in peripheral blood of humanized mice (**Fig. S2**) and the clearance rates of extracellular HBV DNA and antigens were fixed as previously estimated values, that is, *μ* = 16.1 d^-1^ [32] and *σ* = 1.00 d^-1^ [33]. Regardless of the proportion, we showed that the model well-captured the experimental quantification data (i.e., extracellular HBV DNA, HBcrAg, HBeAg, and HBsAg) over time with best fit parameters (**Fig. 3AB** and **Fig. S3**). However, comparing SSR (SSR = 19.9, 23.1, 31.9 for *x* = 0, 0.5, 0.8, respectively), we found a model considering that cccDNA is the dominant source of HBsAg (i.e., *x* = 0) best describes our data. In the following, we discussed the parameter values of the multiscale mathematical model with no HBsAg production from iDNA (see **Discussion**). The estimated parameters and fixed initial values are listed in **Table 2, Table S1** and **Table S2**, respectively.

**Table 2.**
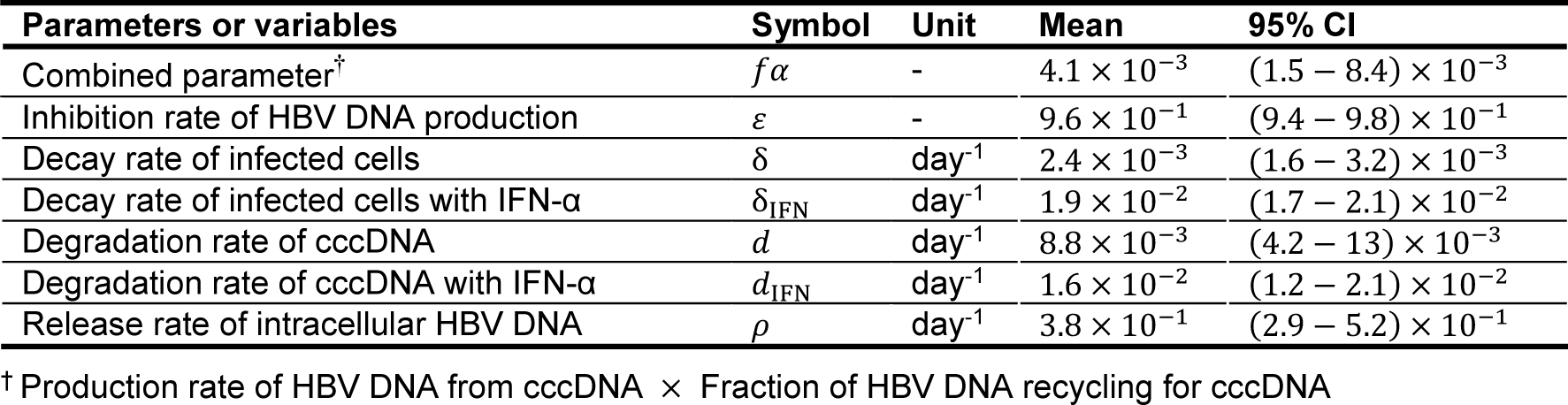
Estimated parameters for HBV infection in humanized mouse.

**Figure 3.**
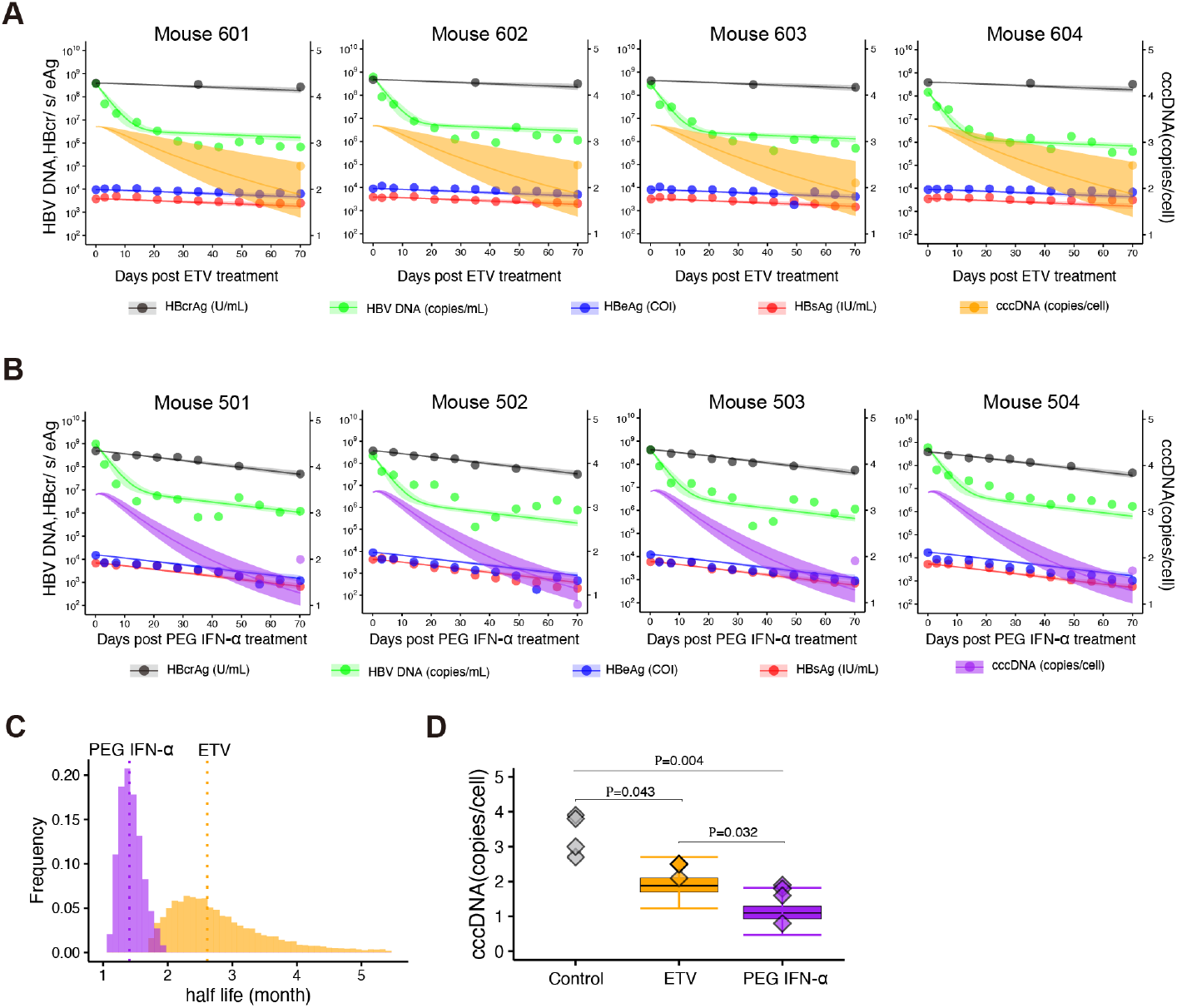
Dynamics of HBV infection in humanized mice: **(A)** and **(B)** show fits of the mathematical model to the extracellular viral markers in peripheral blood of humanized mice treated with ETV or PEG IFN-α (black: HBcrAg, green: HBV DNA, blue: HBeAg, red: HBsAg). The shaded regions correspond to 95% posterior intervals and the solid curves give the best-fit solution (mean) for Eqs. (S17-20) or (S30-33) to the time-course dataset. All data were fitted simultaneously. The intrahepatic cccDNA levels (dots) measured in liver samples under treatment with ETV or PEG IFN-α and its model predictions by Eq.(S22) or (S35) (lines and shaded regions are the mean and 95% posterior intervals) are also shown in orange and purple with the different scale and unit of y-axis, respectively. **(C)** The distribution of the half-life of cccDNA, log 2 /*d*, under treatment with PEG IFN-α inferred by MCMC computations. **(D)** Comparisons of predicted cccDNA copies/cell by Eq.(S22) or (S35) with estimated parameters and the measured cccDNA levels at before (i.e., baseline) and 70 days after ETV or PEG IFN-α treatment in humanized mice. Black line indicates the median, box and whiskers show the interquartile range (IQR) and 1.5×IQR, respectively.

### Predicting intrahepatic cccDNA dynamics from extracellular viral makers

When we applied the multiscale mathematical model to the evaluation of the drug effects on viral replication and amount of cccDNA, it was assumed that ETV almost completely blocks intracellular HBV replication and *de novo* infections (i.e., potent antiviral effect) but has no direct effect on cccDNA degradation, as reported previously (**Supplementary Note 2**) [22, 34-36]. We found the mean half-life of cccDNA was 109 days in the humanized mice under ETV treatment (**Fig. 3C** and **Table 2**). In addition to the potent antiviral effect of PEG IFN-α (i.e., no *de novo* infections, see **Supplementary Note 3**) as other reports [37], our analysis demonstrated that PEG IFN-α treatment significantly reduces the half-life of cccDNA to around 49 days (**Fig. 3C** and **Table 2**), implying PEG IFN-α promotes cccDNA degradation. This calculation is supported by our previous mouse experiments showing that PEG IFN-α treatment for 42 days reduced cccDNA levels to 23-33%, which was semi-quantified with the bands detected by southern blot [38] (**Table S3**). Note that this cccDNA half-life upon antiviral treatment is estimated under the assumption that no *de novo* infections occur owing to the robust antiviral effects; the cccDNA half-life value can be even shorter when low-level *de novo* infections occur during ETV and/or PEG IFN-α treatment (**Supplementary Note 2**,**3**).

Importantly, the intrahepatic cccDNA levels experimentally measured in liver samples from humanized mice (cccDNA was measured by collecting the liver from sacrificed mice and digesting the tissue with plasmid-safe ATP-dependent deoxyribonuclease DNase [PSAD], followed by absolute quantification by droplet digital PCR [ddPCR]) [38, 39] were confirmed to be within the range of values calculated by our mathematical models, Eq. (S22) and Eq. (S35) in **Supplementary Note 2, 3**, assuming the mean cccDNA level of non-treated humanized mice as its initial value (**Fig. 3D**). Note that the individual cccDNA levels and the corresponding model projections are shown in **Fig. 3AB** with the different scale and unit of y-axis (i.e., linear scale). Taken together, our extended approach with those viral markers in peripheral blood predicted intrahepatic cccDNA dynamics and captured the reduction of the half-life of cccDNA *in vivo* by treatment with PEG IFN-α.

## Discussion

So far, mathematical models with several “compartmentalized stages” of intracellular HBV replication (i.e., described by ODEs which cannot explain the time-dependent extracellular viral marker production, for example) have been proposed [12, 13]. Here, we developed a multiscale mathematical model explicitly including intracellular and intercellular HBV infection, described by age-structured PDEs, for quantifying HBV viral dynamics based on *in vitro* and *in vivo* experimental data. Then, the amount of intrahepatic cccDNA and its dynamics are predicted by quantification of serum viral markers - HBV DNA, HBsAg, HBeAg and HBcrAg - in this multiscale model. Our mathematical model based on results from the humanized mouse model without HBsAg produced from iDNA was well fitted. In other words, the levels of iDNA-derived HBsAg are assumed to be negligible compared with those derived from cccDNA. This is because cccDNA is known to be the major source of HBsAg production in HBeAg-positive patients and animal models [40]. On the other hand, previous papers reported that iDNA of the HBsAg region may contribute to HBsAg production rather than cccDNA in HBeAg-negative patients [5, 6, 40]. When we applied our multiscale model to quantify intrahepatic cccDNA in HBV-infected individuals, it would be necessary to include HBsAg produced from iDNA especially for HBeAg-negative patients.

In this study, we calculated cccDNA copy number and half-life in PHH and humanized mice. Concerning copy number, the number of cccDNA copies was higher in PHH than mouse. Recent studies have reported little amplification of cccDNA copy number in infected cells after primary infection [41]. This suggests that cccDNA copies in infected cells depend on the amount of cccDNA formed at the initial infection, which correlates with the higher amount of cccDNA in PHH, which are exposed to a large amount of HBV at the initial infection, and the lower amount in humanized mice [42, 43]. On the other hand, cccDNA half-life was shorter in PHH than humanized mouse. cccDNA persistence in hepatocytes requires supplementation of cccDNA by intracellular recycling of the viral genome and/or de novo infection [41]. Although viral recycling will occur in PHH and humanized mice, de novo reinfection is rarely observed in PHH. This is because viral infection in PHH requires the addition of PEG8000 [41], which was not added after the initial infection in this experiment. Thus, one of the pathways to maintain cccDNA levels does not work in PHH, resulting in a calculated cccDNA half-life that is shorter than that in humanized mice.

Our study has some limitations as follows. First, the experimental quantification method of cccDNA: We quantified cccDNA by PCR-based methods, because of the requirement for a large number of quantifications for the mathematical model. Standardization of the detection method for cccDNA by real-time PCR has been discussed over the years [38, 39]. We have to be careful about the possible overestimation of cccDNA amount even if minimizing the contamination of rcDNA by PSAD digestion as used in this study. However, the cccDNA half-life value estimated by our method is roughly unaffected by a slight shift of cccDNA levels. We minimized this limitation by comparing the PCR-based cccDNA quantification data with the values detected by Southern blot in HBV-infected chimeric mice (**Fig. 3D, Table 2**, and **Table S3**). Second, our mathematical model has a few assumptions underlying the intracellular and intercellular HBV propagation. We assumed negligible *de novo* infections under ETV and PEG IFN-α treatment because NAs and PEG IFN-α inhibit HBV replication by around 100% (i.e., *ε* ≈ 1) (**Supplementary Note 2**,**3**). The assumption may overestimate the mean half-life of cccDNA. After additional datasets on the time-course of the viral markers with different intensities of NAs and PEG IFN-α treatments become available, the inhibition rate can be determined more precisely and our estimation will be improved. Although the current simple but quantitative mathematical model successfully predicts the amount of cccDNA in humanized mice from our extracellular viral markers, more detailed mathematical modeling that improves these limitations will be beneficial for further precise estimation on cccDNA dynamics.

In summary, our multiscale mathematical model combined with extracellular viral markers - HBsAg, HBcrAg, HBeAg and HBV DNA - predicts the amount of intrahepatic cccDNA *in vivo* and may open new avenues to analyze clinical data for understanding cccDNA dynamics in patients.

## Methods

### HBV infection in primary human hepatocytes

PHH used for the HBV infection assay were maintained according to the manufacturer’s protocol (Phoenix Bio Co., Ltd, Hiroshima, Japan). HBV (genotypeD) used as the inoculum was recovered from the culture supernatant of Hep38.7-Tet cells cultured under tetracycline depletion and concentrated up to 200-fold by polyethylene glycol concentration[44]. PHH were seeded into 96-well plates at 7×10^4^ cells/well and were inoculated with HBV at 8,000 genome equivalents (GEq)/cell in the presence of 4% polyethylene glycol 8,000 (PEG8000) for 16 h. After washing out free HBV, PHH were continuously treated with ETV at 1 μM or were not treated (control). Cell division is known to reduce the cccDNA per cell in HBV-infected cells[26]; therefore, to avoid this, we maintained PHH at 100% confluent conditions during the entire infection assay. Moreover, a high concentration of DMSO was included in the culture medium as described previously[45], which does not allow cell growth and prevents cccDNA loss by cell division[25-27]. Since we had confirmed by cell counting that primary culture of human hepatocytes did not significantly proliferate over one month under the above conditions[45], cell growth dynamics were ignored in our analysis. Culture supernatant from HBV-infected cells and the cells were recovered to quantify HBV DNA in the culture supernatant, total HBV DNA in the cells, and cccDNA by real-time PCR. For real-time PCR, the primer-probe sets used in this study were 5’-AAGGTAGGAGCTGAGCATTCG-3’, 5’-AGGCGGATTTGCTGGCAAAG-3’, and 5’-FAM-AGCCCTCAGGCTCAGGGCATAC-TAMRA-3’ for detecting HBV DNA and 5’-CGTCTGTGCCTTCTCATCTGC-3’, 5’-GCACAGCTTGGAGGCTTGAA-3’, and 5’-CTGTAGGCATAAATTGGT(MGB)-3’ for cccDNA[44].

In the assay shown in **Fig. 1**, a large amount of HBV (8000 GEq/cell) is exposed to PHH on day 0, which is the condition in which about 80% of the PHH is infected; thus, the cccDNA amount is high at day 1. Previous papers also reported that cccDNA is readily detected as early as day 2 after HBV inoculation and remains at a similar level over time[27, 46]. Note that the time “day 1” in our study means 24 h after the end of HBV inoculation (16 h), indicating 40 h after starting HBV inoculation. In addition, HBV infection did not spread because PEG8000, which supports viral attachment on the cell surface[47], was not added to the culture medium after day 1, which resulted in the cccDNA initially forming in the cells without increasing, showing similar amounts of cccDNA on day 1 and day 31.

### HBV infection of humanized mouse

Humanized mice were purchased from Phoenix Bio Co., Ltd. (Hiroshima, Japan). The animal protocol was approved by the Ethics Committees of Phoenix Bio Co., Ltd (Permit Number:2200). These mice were infected with HBV at 1.0 × 10^6^ copies/mouse that was obtained from human hepatocyte chimeric mice previously infected with genotype C2/Ce, as described previously[48]. Day 53 after inoculation, HBV-infected mice, which showed a plateau of HBV levels in serum, were treated with ETV (at a dose of 0.02 mg/kg, once a day) or PEG IFN-α (at a dose of 0.03 mg/kg, twice a week) continuously for over 70 days (**Fig. 3AB** and **Fig. S1B**). The human albumin level in the serum was measured as described previously[49]. The HBV DNA titer was measured by real-time PCR as previously described[50]. HBsAg, HBcrAg, and HBeAg were measured by chemiluminescent enzyme immunoassay using a commercial assay kit (Fujirebio Inc., Tokyo, Japan). The detection limit of the HBsAg assay and HBcrAg assay were 0.005 IU/ml and 1.0 kU/ml, respectively. The cut-off index (COI) of the HBeAg was <1.00 (**Fig. 3AB** and **Fig. S3**). Intrahepatic HBV cccDNA was extracted from a dissected liver treated with PSAD to digest genomic DNA and rcDNA as described previously[51] (**Fig. 3D**). Genomic DNA was isolated from the livers of chimeric mice using the phenol/chloroform method as previously described[52]. The cccDNA-specific primer-probe set for cccDNA amplification was used for ddPCR assay[51]. After the generation of reaction droplets, intrahepatic cccDNA was amplified using a C1000 touch™ Thermal Cycler (Bio-Rad, Hercules, California, USA). In all cases, intrahepatic cccDNA values were normalized by the cell number measured by the hRPP30 copy number variation assay (Bio-Rad, Pleasanton, California, USA)[53]. Of note, hRPP30 levels were separately determined using DNA that was not treated with PSAD. Group means of the difference in cccDNA/hepatocyte were compared by unpaired t-test.

### Data fitting and parameter estimation

#### (1) Data analysis for HBV infection on PHH

We categorized datasets as follows: [condition 1 = No ETV treatment], [condition 2 = ETV treatment from day 0] and [condition 3 = ETV treatment from day 9] (**Fig. S1A**). To assess the variability of kinetic parameters and model predictions, we performed Bayesian inference for the dataset of condition 1, 2 and 3 using Markov chain Monte Carlo (MCMC) sampling[11]. A statistical model adopted from Bayesian inference assumed that measurement error followed a normal distribution with mean zero and constant variance (error variance). Simultaneously, we fitted Eqs. (1-3) and Eqs. (1-2)(4) to the experimental data of intracellular HBV DNA and cccDNA, and extracellular HBV DNA in condition 1 and condition 2, 3, respectively (**Fig. 1B**). Note that we estimated model parameters (i.e., *α, f, d, ρ, d*_*E*_, *ε*) for all conditions as common values because the HBV used in this assay is identical. On the other hand, susceptibility and permissiveness of PHH to HBV are known as heterogeneity; thus, we used different initial values (i.e., *C*(0), *D*(0), *Q*(0)) for each condition (**Table 1**). Distributions of model parameters and initial values were inferred directly by MCMC computations[11].

#### (2) Data analysis for HBV infection on humanized mouse

To quantify HBV infection and the antiviral effect of ETV or IFN-α in humanized mice, we also performed Bayesian inference using MCMC sampling because the inter-individual variations are almost negligible. We here used a previously estimated half-life of extracellular HBV DNA in peripheral blood (PB), that is, 62 minutes (*μ* = 16.1 d^-1^)[32], and that of extracellular HBsAg in PB, 0.69 day (*σ* = 1 d^-1^)[33]. Simultaneously, we fitted Eqs. (*S*17-*S*20) and Eqs. (*S30*-*S*33) to the experimentally measured extracellular HBV DNA, HBcrAg, HBeAg and HBsAg obtained from HBV-infected humanized mice treated with ETV and PEG IFN-α, respectively (**Fig. 3AB** and **Fig. S3**), and estimated *d, d*_*IFN*_ and ρ ρ (**Table 2** and **Table S1**). Note that we fixed all initial values as initial points of our dataset (**Table S2**), and the decay rates of infected cells were separately estimated from h-Alb in PB of the humanized mice (**Fig. S2, Table 2** and **Table S1**).

### Statistical analysis

Mathematical modeling, transformation to the reduced model and its linearization are described in **Supplementary Note 1**,**2**,**3** in detail. All analyses of samples were conducted using custom scripts in R and were visualized using RStudio. For comparisons between groups, Mann-Whitney U tests and t test were used. All tests were declared significant for *p* < 0.01.

## LIST OF SUPPLEMENTARY MATERIALS

**Figure S1** | Summary of HBV infection datasets

**Figure S2** | Experiments using HBV-infected humanized mice

**Figure S3** | Fitting of the mathematical model to the extracellular viral markers in peripheral blood of humanized mice treated with ETV or PEG IFN-α considering HBsAg production from iDNA

**Table S1** | Estimated parameters for HBV infection in humanized mouse considering HBsAg production from iDNA

**Table S2** | Fixed initial values for HBV infection in humanized mouse

**Table S3** | Quantified results for cccDNA in HBV infected mouse

**Supplementary Note 1** | Transformation to a system of ODEs from a PDE multiscale model

**Supplementary Note 2** | Linearized equations under potent NAs treatment in humanized mouse

**Supplementary Note 3** | Linearized equations under potent PEG IFN-α treatment in humanized mouse

## AUTHOR CONTRIBUTIONS

KW, SI and YT designed the research. MI, SH, ST, LA, MD, KW and YT conducted the experiments. KSK, HP, SN, ASP and SI carried out the computational analysis. KW, SI, and YT supervised the project. All authors contributed to writing the manuscript.

## ACKNOWLEDGMENTS

This study was supported in part by a Grant-in-Aid for JSPS Research Fellows 20J00868 (to M.I.), 21K15453 (to M.I.); Scientific Research (KAKENHI) B 18H01139 (to S.I.), 16H04845 (to S.I.), 20H03499 (to K.W.), 21H02449 (to K.W.); Scientific Research in Innovative Areas 20H05042 (to S.I.); the Ministry of Education, Culture, Sports, Science, and Technology, 20K16996 (to S.H.); AMED Strategic International Brain Science Research Promotion Program 22wm0425011s0302 (to K.A.); AMED JP22dm0307009 (to K.A.); AMED CREST 19gm1310002 (to S.I.); AMED Development of Vaccines for the Novel Coronavirus Disease, 21nf0101638s0201 (to S.I.); AMED Japan Program for Infectious Diseases Research and Infrastructure, 22wm0325007s8002 (to S.H.), 22wm0325007j0103 (to K.W.), 22wm0325007h0001 (to S.I.), 22wm0325004s0201 (to S.I.), 22wm0325012s0301 (to S.I.), 22wm0325015s0301 (to S.I.); AMED Research Program on Emerging and Re-emerging Infectious Diseases 21wm0325007s8002 (to S.H.), 22fk0108140s0802 (to S.I.); AMED Research Program on HIV/AIDS 22fk0410052s0401 (to S.I.); AMED Program for Basic and Clinical Research on Hepatitis 22fk0210094 (to S.I.); AMED Program on the Innovative Development and the Application of New Drugs for Hepatitis B 22fk0310504j0001 (to K.W.), 22fk0310504h0501 (to S.I.); AMED International Collaborative Research Program Strategic International Collaborative Research Program (SICORP) 22jm0210068j0004 (to K.W.); AMED Research Program on Hepatitis 19fk0210036h0502 (to S.I.), 19fk0210036j0002 (to K.W.), 19fk0310114h0103 (to S.I.), 19fk0310114j0003 (to K.W.), 19fk0310101j1003 (to K.W.), 19fk0310103j0203 (to K.W.), JP21fk0310101 (to Y.T.); JST MIRAI JPMJMI22G1 (to S.I. and K.W.); Moonshot R&D JPMJMS2021 (to K.A. and S.I.) and JPMJMS2025 (to S.I.); The National Research Foundation of Korea (NRF) grant funded by the Korea government (MSIT) (2022R1C1C2003637) (to K.S.K.); National Institutes of Health grants R01-OD011095, R01-AI078881, and R01-AI116868 (to A.S.P); Smoking Research Foundation (to K.W.); The Takeda Science Foundation (to K.W.); Taiju Life Social Welfare Foundation (to K.W.); Shin-Nihon of Advanced Medical Research (to S.I.); SECOM Science and Technology Foundation (to S.I.); The Japan Prize Foundation (to S.I.).

## Supplementary Information

**Figure S1.**
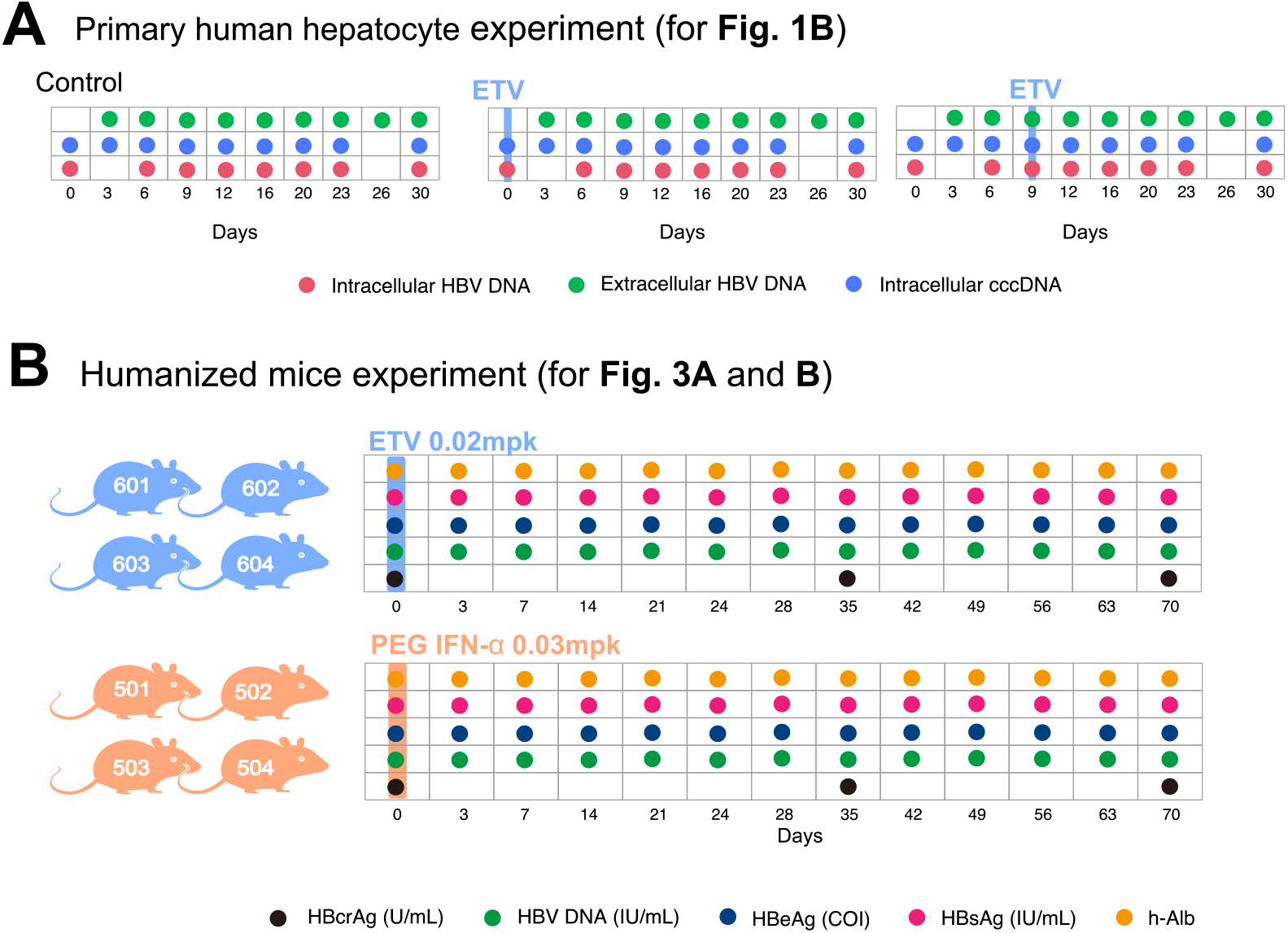
Summary of HBV infection datasets: Detailed data-sampling schedule for HBV-infected **(A)** primary human hepatocytes, and **(B)** humanized mice.

**Figure S2.**
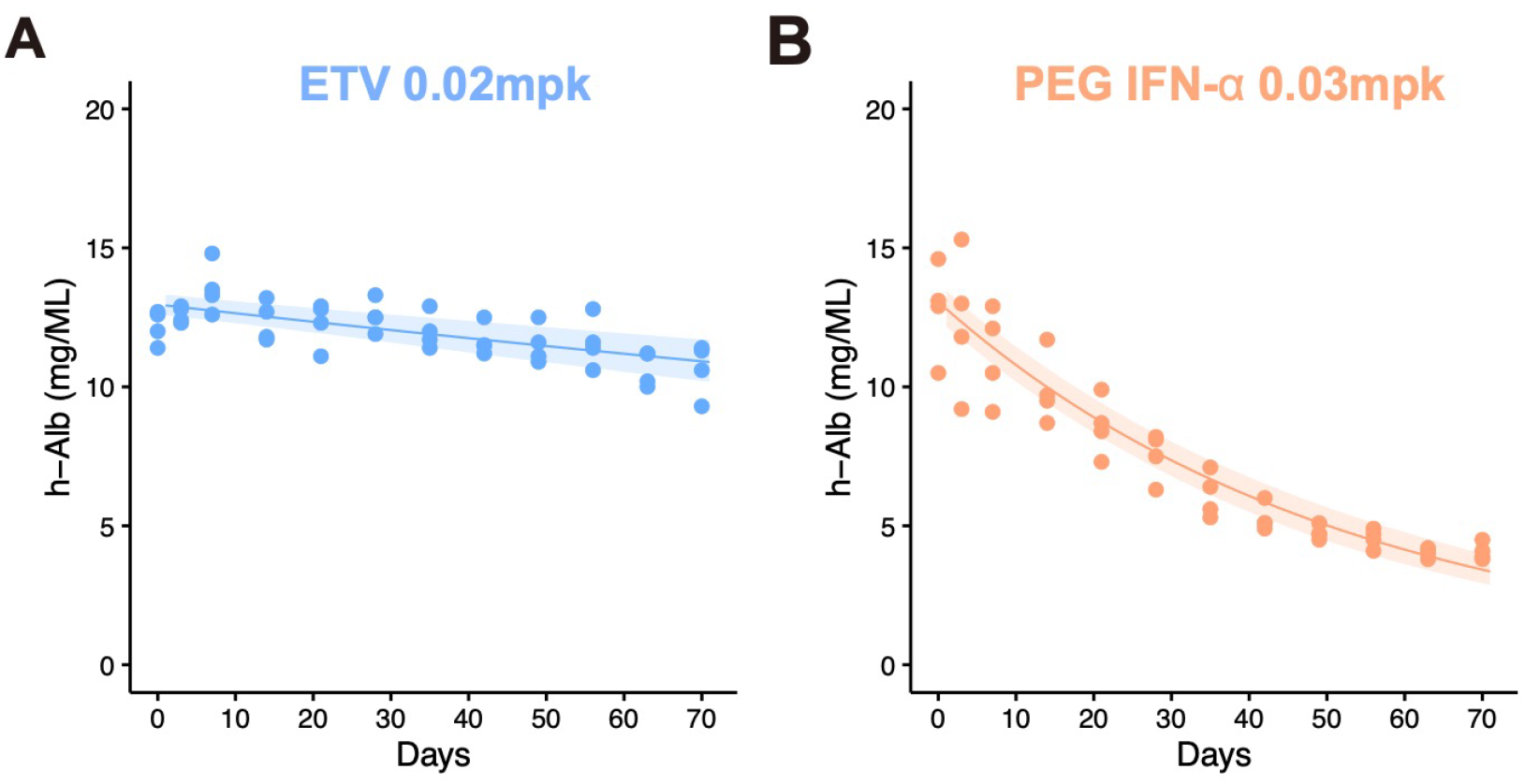
Experiments using HBV-infected humanized mice: Decay characteristics for h-Alb in peripheral blood of humanized mice treated with **(A)** ETV or **(B)** PEG IFN-α. The shaded regions correspond to 95% confidence intervals and the solid curves give the best-fit solution (mean) for a single decay model to the time-course dataset.

**Figure S3.**
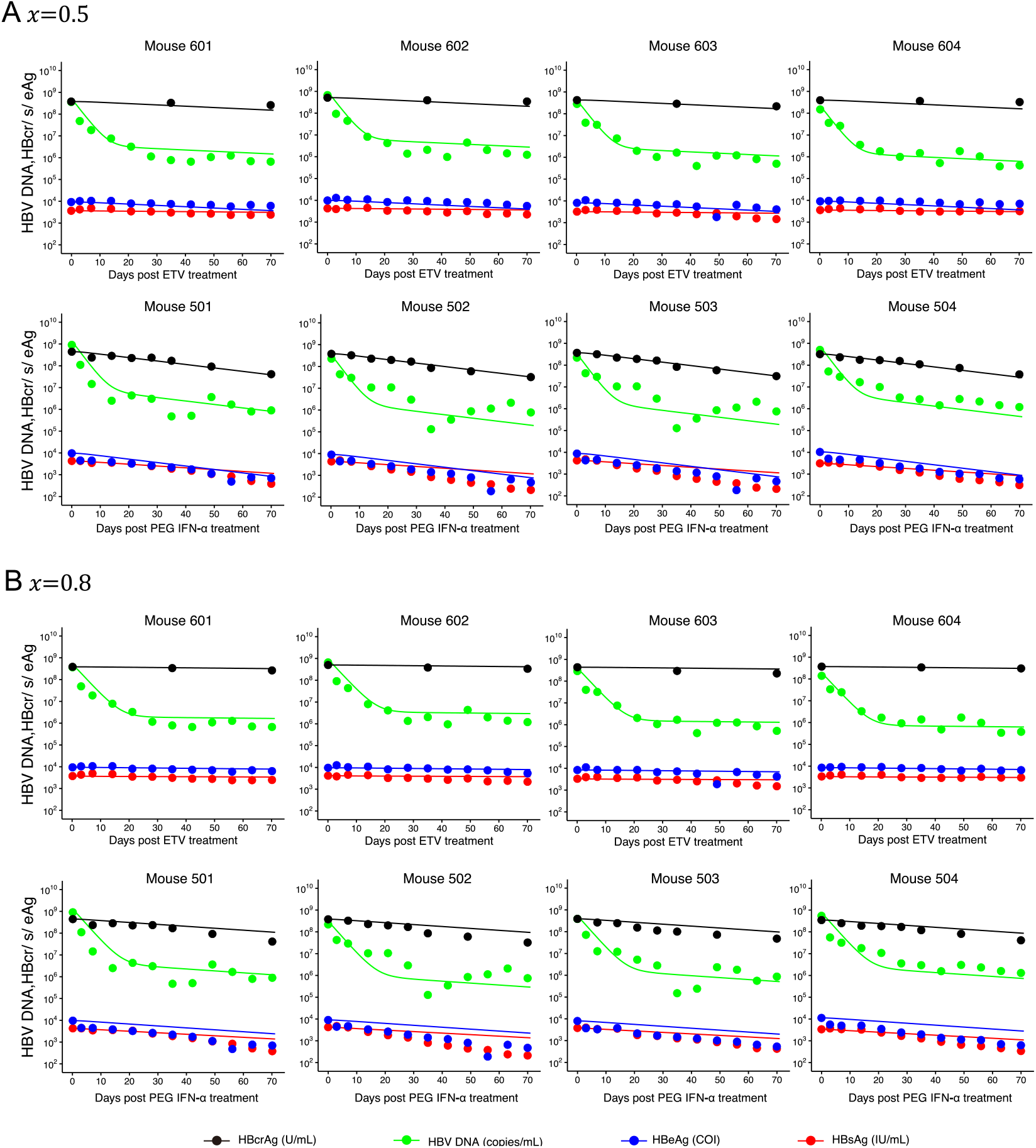
Dynamics of viral makers in HBV infected humanized mice: Fitting of the mathematical model to the extracellular viral markers in peripheral blood of humanized mice treated with ETV or PEG IFN-α considering HBsAg production from iDNA (*x* = 0.5 or 0.8).

**Table S1.** Estimated parameters for HBV infection in humanized mouse considering HBsAg production from iDNA.

| Parameters or variables | Symbol | Unit | Value |
| --- | --- | --- | --- |
| **$x = 0.5$** |  |  |  |
| Combined parameter<sup>†</sup> | $f\alpha$ | - | $5.4 \times 10^{-3}$ |
| Inhibition rate of HBV DNA production | $\varepsilon$ | - | $9.7 \times 10^{-1}$ |
| Decay rate of infected cells | $\delta$ | day<sup>-1</sup> | $2.4 \times 10^{-3}$ |
| Decay rate of infected cells with IFN-α | $\delta_{\text{IFN}}$ | day<sup>-1</sup> | $1.9 \times 10^{-2}$ |
| Degradation rate of cccDNA | $d$ | day<sup>-1</sup> | $1.2 \times 10^{-2}$ |
| Degradation rate of cccDNA with IFN-α | $d_{\text{IFN}}$ | day<sup>-1</sup> | $1.7 \times 10^{-2}$ |
| Release rate of intracellular HBV DNA | $\rho$ | day<sup>-1</sup> | $3.8 \times 10^{-1}$ |
| Residual sum of squares | -- | - | 23.123 |
| **$x = 0.8$** |  |  |  |
| Combined parameter<sup>†</sup> | $f\alpha$ | - | $1.3 \times 10^{-2}$ |
| Inhibition rate of HBV DNA production | $\varepsilon$ | - | $9.9 \times 10^{-1}$ |
| Decay rate of infected cells | $\delta$ | day<sup>-1</sup> | $2.4 \times 10^{-3}$ |
| Decay rate of infected cells with IFN-α | $\delta_{\text{IFN}}$ | day<sup>-1</sup> | $1.9 \times 10^{-2}$ |
| Degradation rate of cccDNA | $d$ | day<sup>-1</sup> | $4.6 \times 10^{-4}$ |
| Degradation rate of cccDNA with IFN-α | $d_{\text{IFN}}$ | day<sup>-1</sup> | $1.2 \times 10^{-3}$ |
| Release rate of intracellular HBV DNA | $\rho$ | day<sup>-1</sup> | $3.2 \times 10^{-1}$ |
| Residual sum of squares | -- | - | 31.947 |

<sup>†</sup> Production rate of HBV DNA from cccDNA × Fraction of HBV DNA recycling for cccDNA

**Table S2.** Fixed initial values for HBV infection in humanized mouse.

| Variable | Symbol | Unit | Value |
| --- | --- | --- | --- |
| **ETV** |  |  |  |
| Initial value for extracellular HBV DNA for Mouse 601 | $V(0)$ | copies/ml | $3.68 \times 10^9$ |
| Initial value for extracellular HBsAg for Mouse 601 | $S(0)$ | IU/ml | $3.75 \times 10^3$ |
| Initial value for extracellular HBeAg for Mouse 601 | $E(0)$ | COI | $9.41 \times 10^3$ |
| Initial value for extracellular HBcrAg for Mouse 601 | $R(0)$ | U/ml | $3.85 \times 10^9$ |
| Initial value for extracellular HBV DNA for Mouse 602 | $V(0)$ | copies/ml | $6.53 \times 10^9$ |
| Initial value for extracellular HBsAg for Mouse 602 | $S(0)$ | IU/ml | $4.14 \times 10^3$ |
| Initial value for extracellular HBeAg for Mouse 602 | $E(0)$ | COI | $9.52 \times 10^3$ |
| Initial value for extracellular HBcrAg for Mouse 602 | $R(0)$ | U/ml | $4.97 \times 10^9$ |
| Initial value for extracellular HBV DNA for Mouse 603 | $V(0)$ | copies/ml | $2.82 \times 10^9$ |
| Initial value for extracellular HBsAg for Mouse 603 | $S(0)$ | IU/ml | $3.22 \times 10^3$ |
| Initial value for extracellular HBeAg for Mouse 603 | $E(0)$ | COI | $8.13 \times 10^3$ |
| Initial value for extracellular HBcrAg for Mouse 603 | $R(0)$ | U/ml | $4.25 \times 10^9$ |
| Initial value for extracellular HBV DNA for Mouse 604 | $V(0)$ | copies/ml | $1.48 \times 10^9$ |
| Initial value for extracellular HBsAg for Mouse 604 | $S(0)$ | IU/ml | $3.56 \times 10^3$ |
| Initial value for extracellular HBeAg for Mouse 604 | $E(0)$ | COI | $8.99 \times 10^3$ |
| Initial value for extracellular HBcrAg for Mouse 604 | $R(0)$ | U/ml | $3.92 \times 10^9$ |
| **PEG IFN-α** |  |  |  |
| Initial value for extracellular HBV DNA for Mouse 501 | $V(0)$ | copies/ml | $9.26 \times 10^9$ |
| Initial value for extracellular HBsAg for Mouse 501 | $S(0)$ | IU/ml | $4.35 \times 10^3$ |
| Initial value for extracellular HBeAg for Mouse 501 | $E(0)$ | COI | $9.79 \times 10^3$ |
| Initial value for extracellular HBcrAg for Mouse 501 | $R(0)$ | U/ml | $4.49 \times 10^9$ |
| Initial value for extracellular HBV DNA for Mouse 502 | $V(0)$ | copies/ml | $2.29 \times 10^9$ |
| Initial value for extracellular HBsAg for Mouse 502 | $S(0)$ | IU/ml | $4.41 \times 10^3$ |
| Initial value for extracellular HBeAg for Mouse 502 | $E(0)$ | COI | $9.08 \times 10^3$ |
| Initial value for extracellular HBcrAg for Mouse 502 | $R(0)$ | U/ml | $3.81 \times 10^9$ |
| Initial value for extracellular HBV DNA for Mouse 503 | $V(0)$ | copies/ml | $3.66 \times 10^9$ |
| Initial value for extracellular HBsAg for Mouse 503 | $S(0)$ | IU/ml | $3.63 \times 10^3$ |
| Initial value for extracellular HBeAg for Mouse 503 | $E(0)$ | COI | $7.59 \times 10^3$ |
| Initial value for extracellular HBcrAg for Mouse 503 | $R(0)$ | U/ml | $3.69 \times 10^9$ |
| Initial value for extracellular HBV DNA for Mouse 504 | $V(0)$ | copies/ml | $5.03 \times 10^9$ |
| Initial value for extracellular HBsAg for Mouse 504 | $S(0)$ | IU/ml | $3.13 \times 10^3$ |
| Initial value for extracellular HBeAg for Mouse 504 | $E(0)$ | COI | $1.04 \times 10^4$ |
| Initial value for extracellular HBcrAg for Mouse 504 | $R(0)$ | U/ml | $3.22 \times 10^9$ |

**Table S3.** Quantified results for cccDNA in HBV infected mouse.

| **Experimental group A** | **cccDNA<sup>†</sup>**<br>**(band volume)** | **Average**<br>**(band volume)** | **% of control** |
| --- | --- | --- | --- |
| untreated control mouse A1 | $5.11 \times 10^7$ | $4.83 \times 10^7$ | 100 |
| untreated control mouse A2 | $4.55 \times 10^7$ | — | — |
| PEG IFN-α treated mouse A1 | $1.74 \times 10^7$ | $1.60 \times 10^7$ | 33 |
| PEG IFN-α treated mouse A2 | $1.46 \times 10^7$ | — | — |
| **Experimental group B** | **cccDNA**<br>**(band volume)** | **Average**<br>**(band volume)** | **% of control** |
| untreated control mouse B1 | $1.31 \times 10^7$ | $1.13 \times 10^7$ | 100 |
| untreated control mouse B2 | $9.44 \times 10^6$ | — | — |
| PEG IFN-α treated mouse B1 | $3.14 \times 10^6$ | $2.62 \times 10^6$ | 23 |
| PEG IFN-α treated mouse B2 | $2.10 \times 10^6$ | — | — |

<sup>†</sup>cccDNA band volume was quantified from Southern blot data<sup>1</sup>. Briefly, mice infected with HBV at 12 weeks were treated with or without PEG IFN-α for 6 weeks, and then they were sacrificed. cccDNA levels were determined by Southern blot in Epicentre-based DNA extracts without proteinase K after PSD digestion. Experimental group A and B were performed as independent experiments.

### Supplementary Note 1: Transformation to a system of ODEs from a PDE multiscale model

We here introduce a multiscale model using partial differential equations (PDEs) that couple intra-, inter- and extra-cellular virus dynamics for analyzing multiscale experimental data of HBV infection (c.f.<u>^2^</u>) (**Fig. 2**):

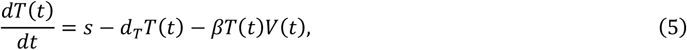

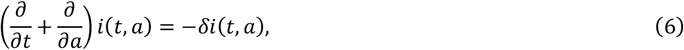

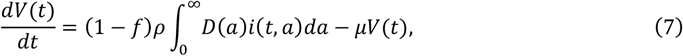

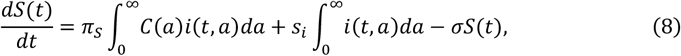

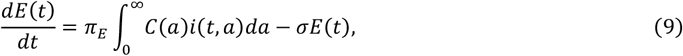

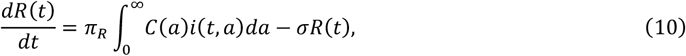

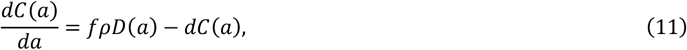

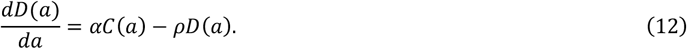

As we recently reported,^<u>3,4</u>^ the multiscale PDE model, Eqs.(5-12), can be transformed into a mathematically identical set of ordinary differential equations as follows. Using the method of characteristics with initial and boundary conditions of *i*(*t, ∂*), we transform Eq. (6) into

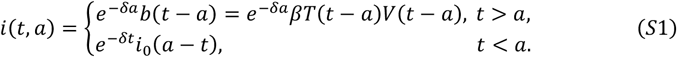

Then, *I*(*t*) is evaluated as follows:

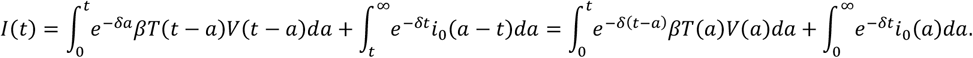

Since 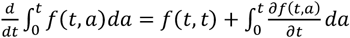, differentiating *I*(*t*) with respect to time *t*, we obtain the following ODE:

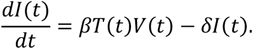

Also, we consider the total amount of cccDNA *CC*(*t*) and the total amount of rcDNA *DD*(*t*), defined by

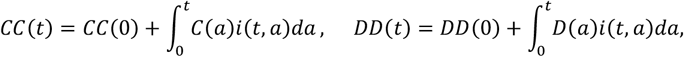

respectively. Then we have

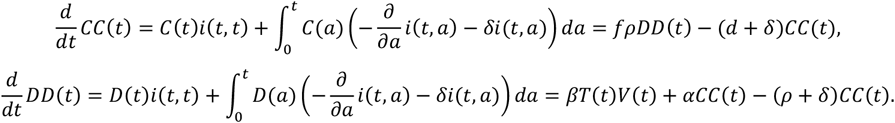

Therefore, the multiscale PDE model is described as the following equivalent system of ODEs:

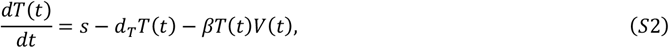

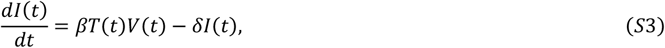

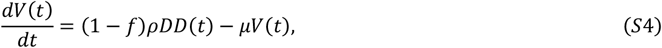

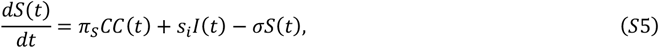

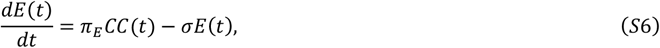

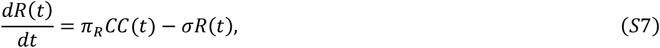

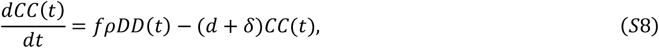

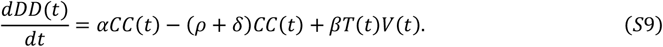

Note that Eqs. (*S*2-*S*9) will be further simplified for the purpose of data analysis depending on the antiviral treatment assumed (see later).

### Supplementary Note 2: Linearized equations under potent NAs treatment in humanized mouse

We assumed that NAs treatment is potent enough that intracellular HBV replications and *de novo* infections are negligible after treatment initiation<u>^5-8^</u>, i.e., the antiviral effectiveness of NAs on intracellular HBV replications is assumed to be 0 < *ε* ≤ 1 and

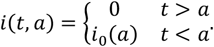

Then Eqs. (*S*2-*S*9) can be simplified as follows:

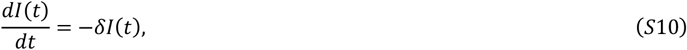

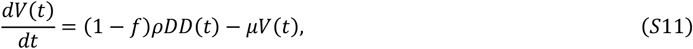

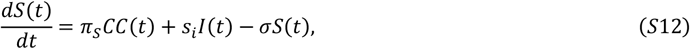

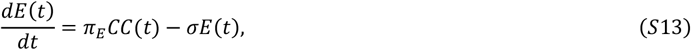

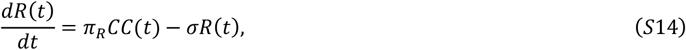

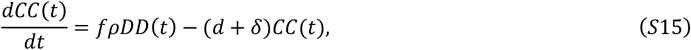

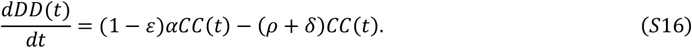

Here we assume that all variables in Eqs. (*S*2-*S*9) are in steady state before treatment initiation<u>^9^</u>, and particularly that the infected cells obtain a stable age distribution, i.e., *i*_0_(*a*) = *βd*(0)*V*(0)*e*^−*δa*^.

Since Eqs. (*S*10-*S*16) are a set of linear ODEs, we directly solve them, and find the following analytical solutions:

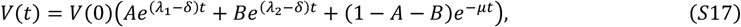

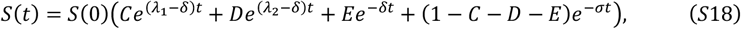

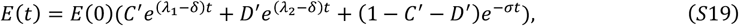

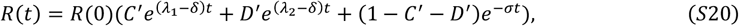

moreover, the total amount of cccDNA *CC*(*t*) and the amount of cccDNA per infected cell 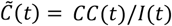 are derived as follows:

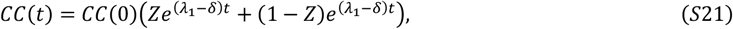

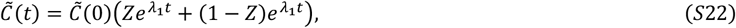

where 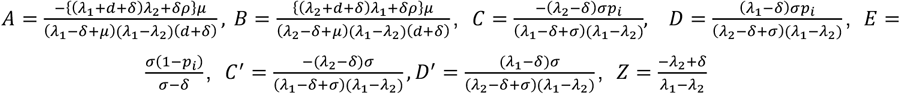 and 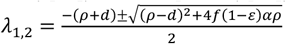. Note that 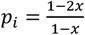 and *x* is the proportion of HBsAg produced from integrated DNA: 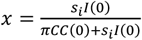.

### Supplementary Note 3: Linearized equations under potent PEG IFN-α treatment in humanized mouse

We also assumed that PEG IFN-α treatment is potent enough that intracellular HBV replication and *de novo* infections are negligible after treatment initiation^6,7,10-12^, i.e., the antiviral effect of PEG IFN-α on intracellular HBV replications is assumed to be 0 < *ε* ≤ 1 and

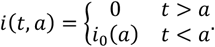

Then Eqs. (*S*2-*S*9) can be simplified to

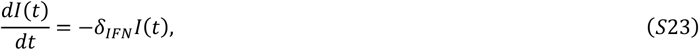

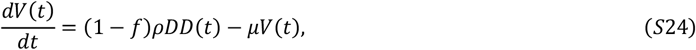

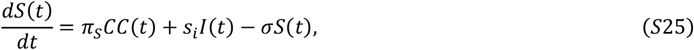

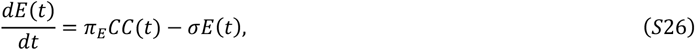

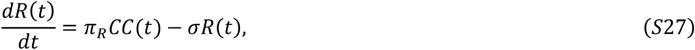

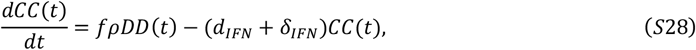

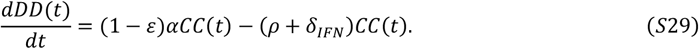

In addition, it has been reported that PEG IFN-α induces interferon-stimulated genes (ISGs) and ISGs potentially degrade intracellular cccDNA. Therefore, we assumed PEG IFN-α increases the cccDNA degradation rate<u>^13^</u>, i.e., *d*_*IFN*_ (> *d*). Similarly, we assume that all variables in Eqs. (*S*2-*S*9) are in steady state before treatment initiation, and that the infected cells have obtained a stable age distribution, i.e., *i*_0_(*a*) = *βd*(0)*V*(0)*e*^−*δa*^. Because PEG IFN-α may enhance the decay rate of infected cells in HBV infection due to cytotoxic effects (but relatively mild), we assumed *δ*_*IFN*_ (≥ *δ*) in the data fitting (**Fig. 3AB** and **Fig. S3**). Solving Eqs. (*S*21-*S*27) we find

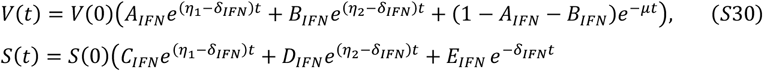

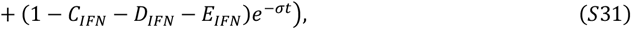

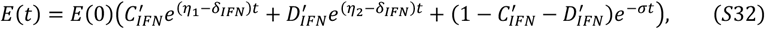

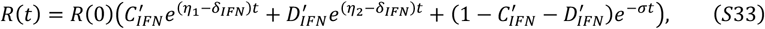

moreover, the total amount of cccDNA *CC*(*t*) and the amount of cccDNA per infected cell 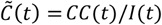 are derived as follows

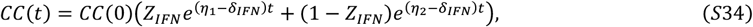

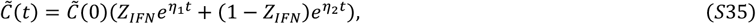

where 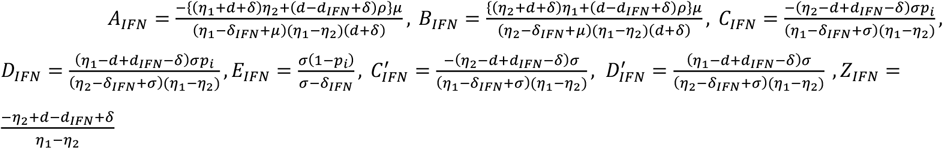 and 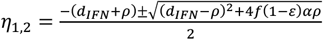. Note that 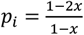 and *x* is the proportion of HBsAg produced from integrated DNA: 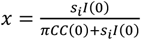.

